# Rete Ovarii Epithelial Cells as an Unappreciated Cell of Origin for Pelvic and Ovarian High-Grade Serous Carcinoma

**DOI:** 10.1101/2025.10.31.685873

**Authors:** Peichao Chen, Xingeng Zhao, Jinpeng Ruan, Xiangmin Lv, Jiyuan Liu, Rui Xu, Kazi Islam, Hongbo Wang, Davie Shi, Madelyn L. Moness, Brianna Carbajal, Gunjan Kumar, Olukemisola A Abioye, Jeniffer McKey, John S Davis, Cheng Wang

## Abstract

Over the past two decades, converging clinicopathologic, molecular, and evolutionary evidence has established that pelvic and ovarian high-grade serous carcinoma (HGSC) originates predominantly from tubal-type epithelia rather than the ovarian surface epithelium. Consequently, Fallopian tube secretory epithelial cells are widely recognized as the principal cell of origin for HGSC. However, the female reproductive tract contains additional tubular epithelial networks whose potential contributions to HGSC pathogenesis remain unexplored. Here, we identify the rete ovarii (RO), a complex network of intra- and extra-ovarian tubules located within the ovarian hilus and mesovarium, as an alternative candidate tissue of origin for HGSC. We demonstrate that genetic and genomic alterations in rete ovarii epithelial cells (ROECs) can drive their malignant transformation, giving rise to tumors that closely recapitulate the histologic and molecular characteristics of human HGSC. Spatially resolved single-cell transcriptomic analyses of tumors derived from *Brca/Trp53/Pten*-deficient ROECs (*BTP*-RO) reveal distinct invasive and immunosuppressive molecular programs. These findings establish ROECs as a previously unrecognized cell of origin for HGSC, expanding the landscape of potential precursor populations beyond the Fallopian tube. The RO-based HGSC model provides a powerful framework for developing origin-informed prevention strategies, early detection approaches, and targeted therapeutics.

## INTRODUCTION

According to the most recent global cancer statistics, an estimated 324,603 women were diagnosed with ovarian cancer and 206,956 women died of the disease in 2022 alone (1). In the United States, the American Cancer Society projects approximately 20,890 new cases and 12,730 deaths from ovarian cancer in 2025 (2). High-grade serous ovarian carcinoma (HGSC) is the most common and lethal subtype of epithelial ovarian cancer (EOC). It is characterized by aggressive clinical behavior, pronounced molecular heterogeneity, and a dismal prognosis, accounting for approximately 70-80% of ovarian cancer-related deaths worldwide(3). Despite the introduction of new therapies, including platinum-taxane chemotherapy, anti-angiogenic agents, and more recently PARP inhibitors, population-level survival gains remain modest (4, 5). The limited therapeutic progress underscores the urgent need for complementary strategies aimed at prevention and early detection to reduce HGSC mortality.

Given that practical prevention can substantially lower disease burden and early-stage disease has excellent outcomes (5-year overall survival for stage I HGSC approaches ∼90%), prevention and early detection represent the most promising approaches to reducing mortality at the population level. Indeed, a substantial fraction of cancers in general are attributable to modifiable risk factors, making prevention a feasible and high-impact strategy (6, 7). The success of tobacco control in reducing lung cancer incidence and the implementation of Pap testing, HPV-based screening, and vaccination in preventing cervical cancer in the United States exemplify the impact of both primary and secondary prevention (2, 8–12). In contrast, efforts to prevent or detect ovarian cancer early have been largely unsuccessful, in part because the etiology of ovarian cancer is incompletely understood. Even the precise tissue(s) of origin for HGSC, which provides the anatomical basis for rational prophylactic strategies, remains a subject of debate (13).

Defining the cell of origin for HGSC is therefore essential for developing effective prevention and early detection strategies. While the molecular landscape of HGSC has been characterized extensively, its histogenetic origin remains uncertain. Historically, the ovarian surface epithelium (OSE) was regarded as the primary source of HGSC (13–16). However, converging clinical and experimental evidence, particularly from BRCA1/2 mutation carriers and genetically engineered mouse models, has shifted the prevailing paradigm toward secretory epithelial cells of the distal fallopian tube as the principal site of origin (17–21), This “tubal origin” hypothesis is supported by the discovery of serous tubal intraepithelial carcinoma (STIC) lesions, which share striking phenotypic, genetic, and developmental similarities with the fallopian tube epithelium (18, 22–25). Nevertheless, the Fallopian tube model does not account for all cases. A considerable fraction of HGSCs is diagnosed without identifiable STIC lesions, even after meticulous pathological review (13, 26), and the population frequency of STIC remains low, including among high-risk BRCA1/2 mutation carriers (27–30). Moreover, the female reproductive tract harbors additional tubular epithelial networks beyond the fallopian tube. These observations, together with emerging studies suggesting additional origins, sustain a long-standing controversy regarding HGSC pathogenesis (13, 31–33).

Here, we provide direct evidence that the rete ovarii (RO), a complex network of intra- and extra-ovarian tubules located in the ovarian hilus and mesovarium(34–36), represents an alternative tubular epithelial source for HGSC. We demonstrate that epithelial cells from RO tubules can undergo malignant transformation, and that the resulting tumors recapitulate the histologic, immunophenotypic, and molecular features of human HGSC. These findings reveal that the RO epithelial cell, an overlooked but persistent component of the ovarian hilus, constitutes a previously unappreciated cell of origin for HGSC.

## RESULTS

### Rete Ovarii Exhibits a Tubular Architecture Resembling the Fallopian Tube

Because HGSC is widely thought to arise from tubular epithelia of the female reproductive tract (37–40), we first examined the gross and microscopic architecture of RO relative to the adjacent fallopian tube (FT). Using a multimodal approach integrating gross imaging, histology, and ultrastructural analysis, we delineated the anatomic and morphologic relationship between the RO and FT. Under a low power stereomicroscope, the adult mouse extraovarian rete (ER) was visible at the junction between the ovarian bursa and peri-ovarian fat pad, appearing as a sac-like structure embedded within the adipose tissue (Fig. 1A). High power stereomicroscopy revealed a network of highly convoluted tubular structures within the sac (Fig. 1B). In neonatal mice, the absence of surrounding fat facilitated isolation of the RO, which appeared as a fine tubular network with a markedly smaller diameter (less than one fifth that of the FT lumen) (Fig. 1C). Scanning electron microscopy (SEM) demonstrated that the RO epithelium comprises both ciliated and non-ciliated (secretory-like) cells, closely resembling the cellular composition of the FT mucosa (Fig. 1D). Low power brightfield imaging further illustrated the anatomical positioning, relative size, and histological organization of the RO in relation to the FT (Fig. 1E). The structural and morphologic convergence observed here supports the hypothesis that the RO epithelium can phenocopy FT features and has the potential to be epithelial source for a subset of tubo-ovarian carcinomas. To corroborate this morphologic similarity at the molecular level, immunohistochemistry for canonical epithelial and lineage markers (Pax8, Foxj1, Ovgp1, He4, and Cdh1, *etc*.) revealed comparable expression patterns between RO and FT epithelia (Fig. S1) (41–44). These findings reveal that RO and FT epithelia share histologic and immunophenotypic features. This anatomic and molecular resemblance highlights a continuum that merits deeper molecular and functional interrogation of the RO as a potential cell of origin for HGSC.

**Figure 1.**
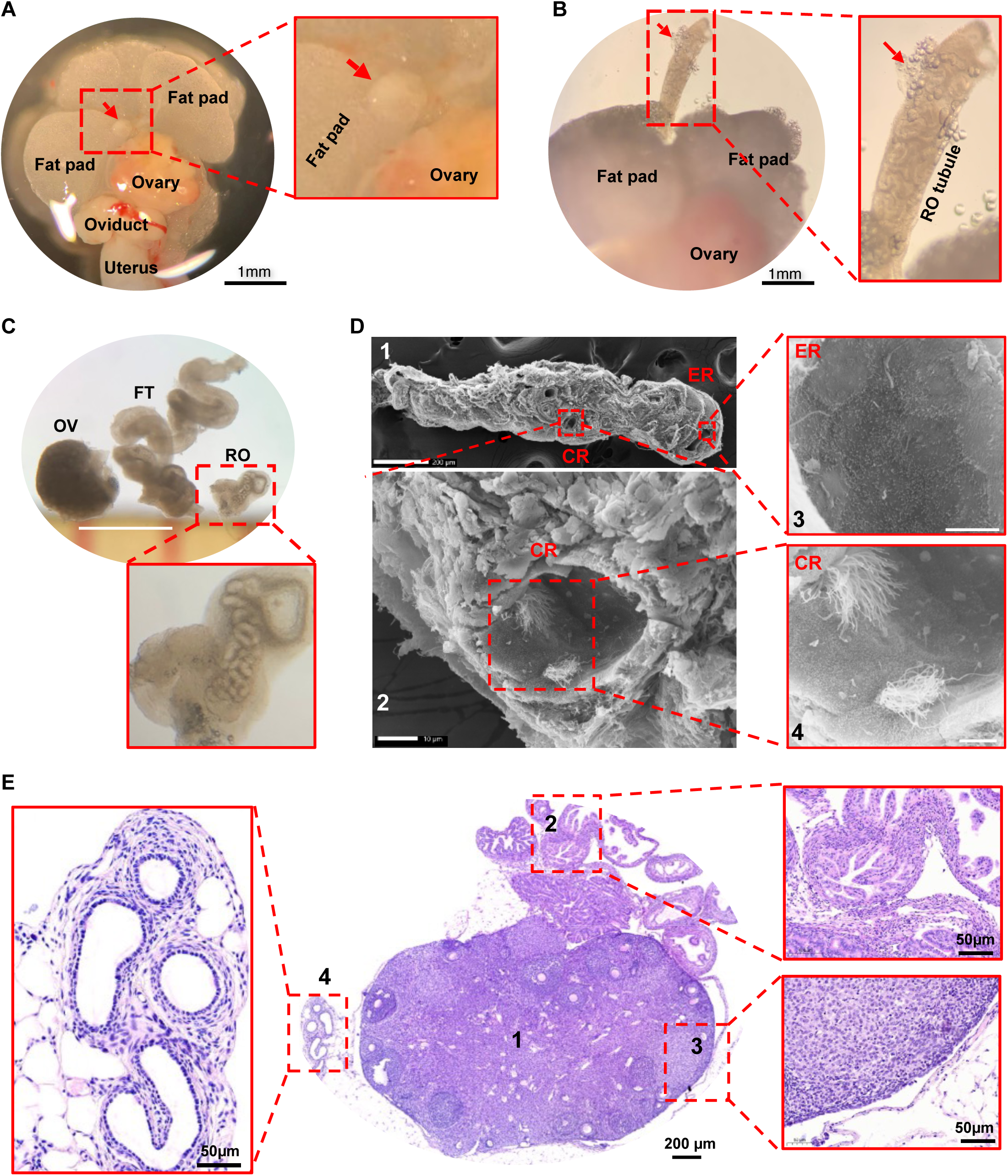
Anatomical and histological characterization of the tubular structure in mouse rete ovarii. A) representative images showing the anatomical location of mouse extraovarian rete ovarii (RO, red arrow), ovary, oviduct, uterus, and peri-ovarian fat pad, providing anatomical context for the RO. Scale bar: 1mm. B) High-resolution images showing the tubular structure (red inset) in an extraovarian RO (red arrow, partially dissected out) from a three-month-old female mouse. The extraovarian RO appears as an extension of contiguous tubular profiles projecting from the ovarian hilum. Scale bar: 1mm C) Relative size comparison of the rete ovarii (RO), ovary (OV), and fallopian tube (FT) in a one-week-old female mouse. A representative image in the lower panel highlights the curved and coiled tubules of a RO from a neonatal female mouse. Scale bar: 1 mm. D) Representative scanning electron microscopy (SEM) images showing the epithelial morphology of RO tubules. D1, RO tubular tissue (×95); scale bar: 200 µm. D2, tightly packed epithelial surface in the connecting fragment of mouse RO (×1400); scale bar: 10 µm. D3, non-ciliated epithelial cells lining the lumen of the extraovarian RO (×5000); scale bar: 5 µm. D4, ciliated epithelium within the collecting RO (×3500). Note apical cilia projecting into the tubule lumen. Scale bar: 5 µm. E) Histological analysis (H&E) showing the relative locations of three epithelial compartments considered candidate cells-of-origin for ovarian carcinogenesis. E1, ovary; scale bar: 200 µm. E2, fallopian tube mucosal epithelium; scale bar: 50 µm. E3, ovarian surface epithelium; scale bar: 50 µm. E4, RO tubule epithelium; scale bar: 50 µm.

### Molecular Signature of the Rete Ovarii and Rete Ovarii Epithelial Cells (ROECs)

To further define the molecular characteristics of RO epithelium and its relationship to the fallopian tube (FT), we established a reference single-cell transcriptomic profile from normal mouse FT tissues and integrated these data with previously published RO single-cell datasets (Fig. 2A, 2B)(45). The integrated analysis resolved 21 distinct cell clusters encompassing both FT and RO lineages (Fig. 2A). Among them, a discrete cluster was enriched for canonical RO markers, including *Amhr2*, *Nr5a1*, and *Fshr*, consistent with prior studies (45). In contrast, the FT secretory epithelial cell (FTSEC) clusters expressed *Ovgp1*, *Ltf*, and *Lcn2*, whereas the ciliated epithelial cell (FTCEC) clusters were enriched for *Ak9*, *Ankfn1*, and *Dnah9*, in agreement with previous profiling of tubal epithelia (46–48) (Fig. 2C).

**Figure 2.**
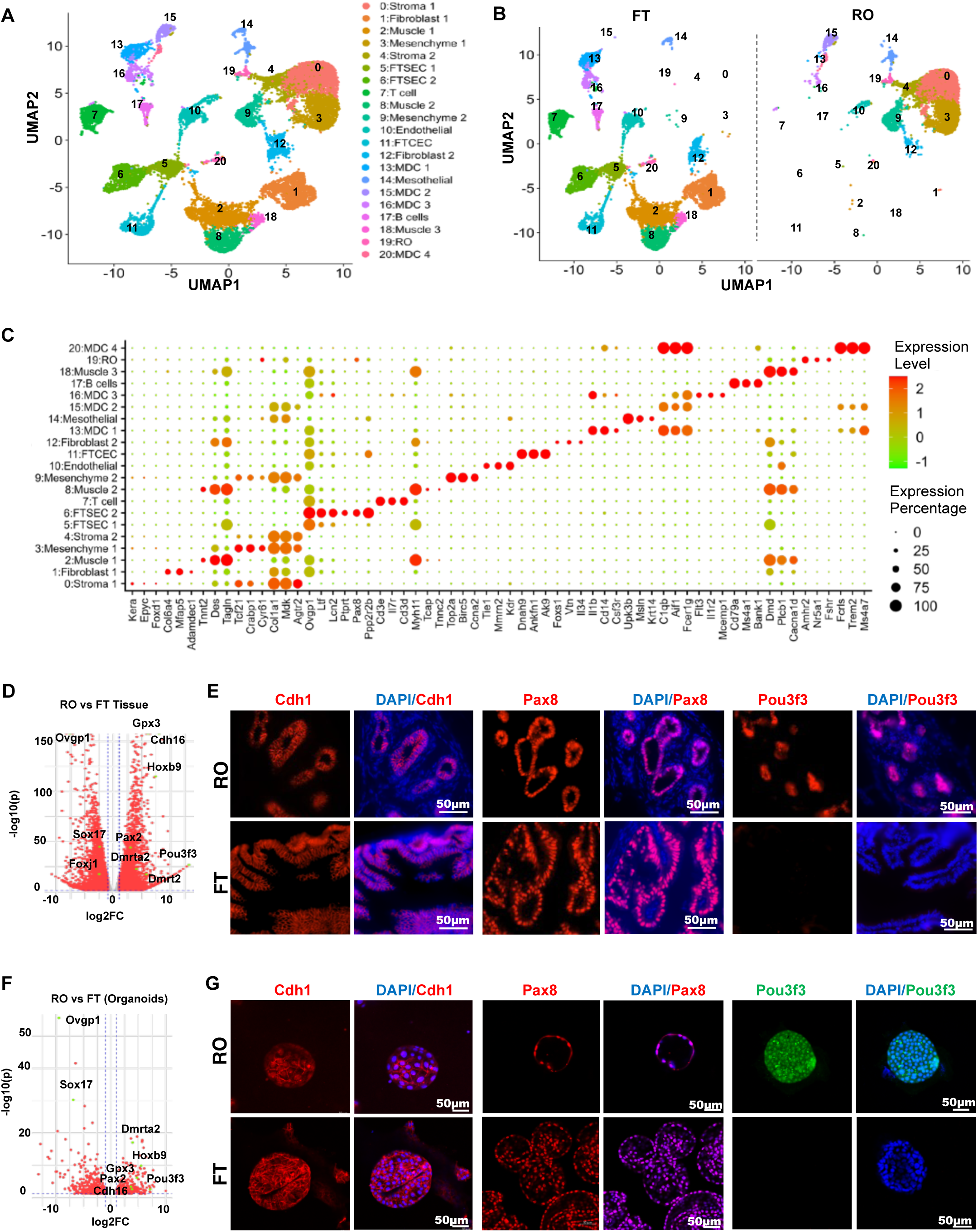
Molecular characterization of RO epithelial cells (ROECs) A) Integrated scRNA-seq UMAP visualization showing 21 major clusters identified in RO and FT tissues by unsupervised clustering. B) UMAP plots illustrating tissue-specific distributions of cell clusters in RO and FT, revealing distinct molecular signatures between epithelial cells from each tissue. C) Dot plot displaying the top three marker genes for each cluster from RO and FT. Dot color represents expression level; dot size indicates the percentage of cells expressing the gene. D) Volcano plot showing genome-wide differentially expressed genes (DEGs) between RO and FT tissues (bulk RNA-seq). DEGs are plotted as log₂ fold change versus −log₁₀(adjusted p). Genes meeting the criteria of adjusted p < 0.05 and |log₂FC| > 1 (n = 3 per group) are displayed, with significant DEGs highlighted in red and key genes labeled in green. E) Fluorescent immunohistochemistry detecting Cdh1 (epithelial marker, red), Pax8 (serous tubular epithelial marker, red), and Pou3f3 (RO epithelial marker, red) in FT and RO tissues. Nuclei stained with DAPI. Scale bar: 50 µm. F) Volcano plot showing DEGs between RO and FT epithelial organoids (bulk RNA-seq). DEGs plotted as log₂ fold change versus −log₁₀(adjusted p), with significant DEGs (adjusted p < 0.05; |log₂FC| > 1; n = 3 per group) highlighted in red; key DEGs in green. G) Fluorescent immunohistochemistry detecting expression of Cdh1 (epithelial cell marker, in red), Pax8 (serous tubular epithelial cell marker, in red), and Pou3f3 (RO epithelial cell marker, in green) in FT and RO organoids. Nuclei were stained with DAPI (blue). Scale bar: 50 µm.

Sub-clustering of RO epithelial cells further revealed two transcriptionally distinct subtypes: one with high expression of *Fshr, Rhox8, Nr5a2,* and *Cyp19a1*, characteristic of secretory-like phenotype; another one with high expression of *Hoxb9*, *Peg3*, *Cyr61*, and *Wnt6*, indicative of ciliated cells (Fig. S2). To validate these transcriptomic findings and explore functional properties unique to RO epithelial cells (ROECs), we established three-dimensional (3D) organoid cultures derived directly from RO tubules (Fig. 2G). Bulk RNA sequencing of RO- and FT-derived tissues and organoids demonstrated distinct lineage-specific transcriptional programs (Fig. 2D, 2F). Among the most differentially expressed genes, *Hoxb9* and *Pou3f3* emerged as robust and specific molecular markers distinguishing ROECs from FT epithelial cells, a finding validated by immunohistochemistry (Fig. 2D-2G, Fig. S1).

Collectively, these data define a distinct molecular signature of the RO epithelium, supporting the notion that the RO represents a tubal-type epithelial derivative with unique transcriptional identity. This molecular resemblance to the FT, combined with its distinct lineage characteristics, underscores the plausibility of the RO as an alternative cell-of-origin for high-grade serous ovarian carcinoma (HGSC).

### Genetic Transformation of RO Epithelial Cells Leads to Aggressive High-Grade Tumors

To investigate the tumorigenic potential of rete ovarii epithelial cells (ROECs), we employed a systematic genetic modification strategy using organoid-based models. A series of ROEC-derived organoids were generated, including wild-type controls (Ctrl), Trp53 knockout (T), *Brca*/*Trp53* double knockout (B;T), B;T with Myc overexpression (B;T;M), B;T with YAP1 overexpression (B;T;Y), Brca knockout/HPV E6/E7 and YAP1 oncogene expression (B;H;Y), and B;T with *Pten* knockout (B;T;P). These genetically engineered organoid lines were subcutaneously implanted into immunodeficient NSG mice to evaluate their tumorigenic capacity. ROECs demonstrated susceptibility to malignant transformation following inactivation of key tumor suppressor pathways and activation of oncogenic signals (Fig. 3A, 3B). For example, tumors derived from B;H;Y-ROECs exhibited histologic features characteristic of high-grade carcinomas, including papillary and solid growth patterns, pleomorphic nuclei, and high proliferative activity as indicated by Ki67 staining (Fig. 3B, 3C).

**Figure 3.**
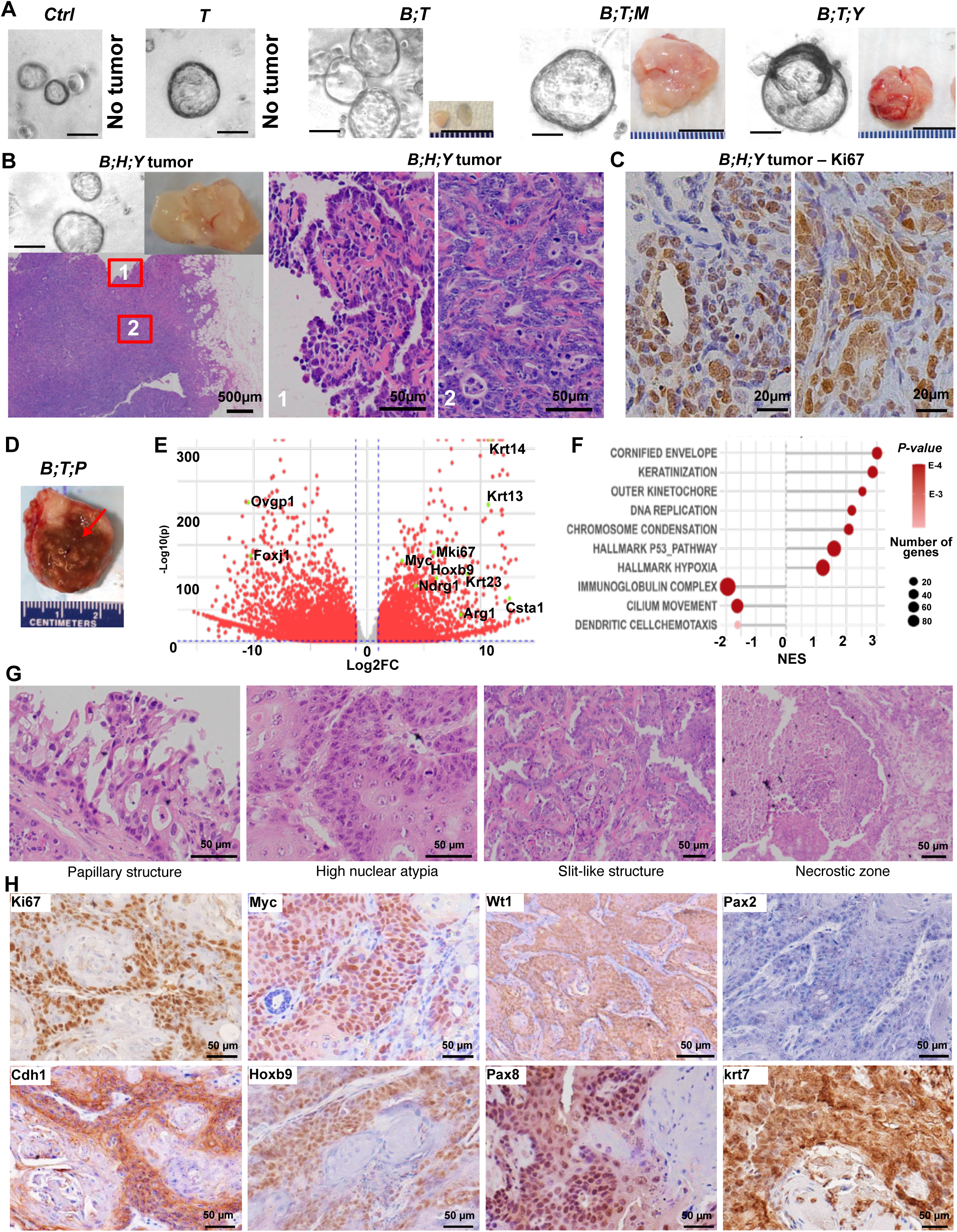
Malignant transformation of RO epithelial cells (ROECs). A) Representative images showing organoids and tumors derived from genetically modified ROECs. Malignant transformation of ROECs was determined by examining their tumorigenic capability in NSG mice. *Ctrl*: wild-type ROECs as control. *T*: ROECs with *Trp53* knockout. *B;T*: ROECs with *Trp53 & Brca* knockout. *B;T;M*: ROECs with inactivation of *Brca & Trp53* and & overexpression of c-MYC. *B;T;Y*: ROECs with inactivation of *Brca & Trp53* and & overexpression of *YAP1*. Please note that no tumor developed in normal ROECs (*Ctrl)* and ROECS with *Tp53* knockout (*T*) groups. *Sample sizes:* n=8 mice per group. *Scale bars:* organoids, 100 µm; tumors, 1 cm. B) Representative images showing histology of tumors derived from *B;H;Y*-ROECs (*ROECs* with *Brca* deficiency + *YAP1* and HPV16 E6/E7 overexpression). Tumorigenic capacity of *B;H;Y-ROECs* was tested in NSG mice (n=8). Representative images of organoids (top left; *scale bar:* 100 µm) and tumor (top right; *scale bar:* 500 µm) derived from *B;H;Y-ROECs* are also presented. High-resolution images on the right show the papillary architecture in insert 1 and solid growth of high-grade tumor cells in insert 2. S*cale bar:* 50 µm. C) Immunohistochemistry showing the expression of Ki67 in tumors derived from *B;H;Y-ROECs implanted to NSG mice. scale bar:* 20 µm. D) A representative image showing tumors derived from *BTP-ROECs* (ROECs with deficiency of *BRCA, TP53,* and *PTEN* genes). These tumors (hereinafter referred to as *BTP*-RO tumor) were generated by intraperitoneally (IP) implanting *BTP-ROECs into* immunocompetent mice. Please note the rapid growth-induced necrosis in these tumors (arrow). E) Volcano plot showing differentially expressed genes (DEGs) in *BTP*-RO tumors when compared to normal RO control tissues (n = 4 per group). DEGs derived from the Bulk RNA-seq data are plotted as log₂ fold change versus - log₁₀(adjusted p). Genes meeting the criteria of adjusted p < 0.05 and |log₂FC| > 1 (n = 4 per group) are displayed. Signature genes of the control tissues and *BTP*-RO tumors are highlighted in green. F) Genes and Pathways enriched in *BTP*-RO tumor. Gene set enrichment analysis (GSEA) based on Bulk RNA-seq data and GO (Gene Ontology) and Hallmark pathway datasets demonstrate that *BTP*-RO tumors are characterized by enrichment of genes and pathways involved in Cornified envelope (GO:0001533), Keratinization (GO:0031424), Outer kinetochore (GO:0000940), DNA replication (GO:0006270), Chromosome condensation (GO:0030261), Immunoglobulin complex (GO:0019814), and Dendritic chemotaxis (GO:0002408). G) H&E staining showing high intra-tumor histological heterogeneity of *BTP*-RO carcinoma. From left to right, papillary growth, solid growth with pronounced nuclear atypia, solid growth with slit-like architecture, and extensive late-stage necrosis. *Scale bar:* 50 µm. H) Immunohistochemistry for epithelial and lineage markers in RO carcinoma (Pax8, Myc, Wt1, Cdh1, Ki67, Hoxb9, and Krt7). *Scale bar:* 50 µm.

We next focused on the *B;T;P*-ROEC model, in which *Brca*, *Trp53*, and *Pten* in ROECs were simultaneously deleted (hereinafter referred to as *BTP-ROEC*). This combination was selected because *Brca*/*Trp53*/*Pten* loss in Fallopian tube epithelium is known to generate tumors that closely resemble human HGSC both histologically and molecularly (21). When *BTP*-ROECs were implanted intraperitoneally into immunocompetent mice, they formed rapidly growing pelvic tumors (hereinafter referred to as *BTP*-RO tumor) that developed necrotic cores filled with serous fluid, consistent with an aggressive, high-grade phenotype rather than a slow-growing low-grade serous carcinoma (Fig. 3D). Bulk RNA sequencing of *BTP*-ROEC-derived tumors revealed extensive transcriptional reprogramming, with upregulation of HGSC signature genes and enrichment of pathways related to hypoxia and keratinization (Fig. 3E). Gene set enrichment analysis (GSEA) confirmed significant activation of hypoxia response, keratinization, and immunosuppressive signaling (Fig. 3F). These results are consistent with prior reports linking hypoxia to immune evasion and metastasis, and keratinization to tumor necrosis in ovarian cancers (49–53). Histologically, *BTP*-RO tumors displayed hallmark features of HGSC, including papillary and solid growth, slit-like glandular structures, pronounced nuclear atypia, and extensive necrosis (Fig. 3G). Immunohistochemistry confirmed strong expression of established HGSC markers such as Pax8, Pcna, Wt1, Cdh1, and Ki67 (3, 21, 54, 55), along with retention of the ROEC marker *Hoxb9* (Fig. 3H). Collectively, these findings demonstrate that Brca/Trp53/Pten-inactivated ROECs give rise to tumors that phenocopy human HGSC.

Notably, hypoxia-related gene expression was elevated even in non-necrotic regions of *BTP*-RO tumors, suggesting spatially heterogeneous hypoxic signaling and possible metabolic or immune interactions within the tumor microenvironment. Because these features could not be resolved by bulk transcriptomic approaches, we next employed spatial transcriptomics to achieve a spatially resolved molecular dissection of *BTP*-RO tumorigenesis.

### Spatial Transcriptomic Profiling Confirms the HGSC Molecular Features of BTP-RO Tumor

To elucidate the molecular and spatial architecture of *BTP*-RO tumors, we performed spatial transcriptomic profiling on tumor tissues and normal RO tubes (as controls) (Fig. 4A). Seventeen distinct spatial clusters were identified across tumor and normal tissues based on genome-wide transcriptional signatures, demonstrating high intratumor heterogeneity and complex tumor– stroma–immune interactions (Fig. 4B-4D). Normal RO tubes segregated into epithelial and stromal clusters and expressed tubal epithelial cell biomarkers (Fig. S3), whereas the *BTP*-RO tumors contained multiple carcinoma sub-clusters, cancer-associated fibroblasts, macrophages, dendritic cells, and B-cell populations (Fig. 4C-4D). The presence of necrotic cores and rapidly proliferating peri-necrotic zones was associated with aggressive growth, indicating that necrosis correlates with tumor progression and metastatic potential. Consistent with the histological classification, canonical high-grade serous carcinoma (HGSC) biomarkers, including *Muc16* (CA125), *Wfdc2* (He4), *Cdh6*, *Pax8*, *Cldn4*, *Epcam*, and *Mki67*, were robustly expressed in carcinoma clusters, whereas the low-grade carcinoma marker Pax2 was absent (Fig.3H and Fig. 4E) (44, 56–61), confirming the HGSC identity of the *BTP*-RO tumor. Expression of Hoxb9 further supported RO epithelial origin of the carcinoma (Fig. 4E). Subtype classification using TCGA-based gene panels revealed very high mesenchymal subtype scores for the mesenchymal subtype of HGSC throughout the tumor (62–65), indicating that high-grade carcinoma derived from transformed *BTP*-ROECs could be classified as mesenchymal subtype of human HGSC (Fig. 4F). Elevated immunoreactive subtype signal in necrotic core highlights the strong spatial heterogeneity and dynamic immune microenvironment within these tumors. Together, these spatial transcriptomic analyses confirm the HGSC molecular identity and extreme intratumor heterogeneity of the *BTP*-RO tumor and underscore the value of this model for investigating tumor evolution, immune suppression, and therapeutic vulnerabilities.

**Figure 4.**
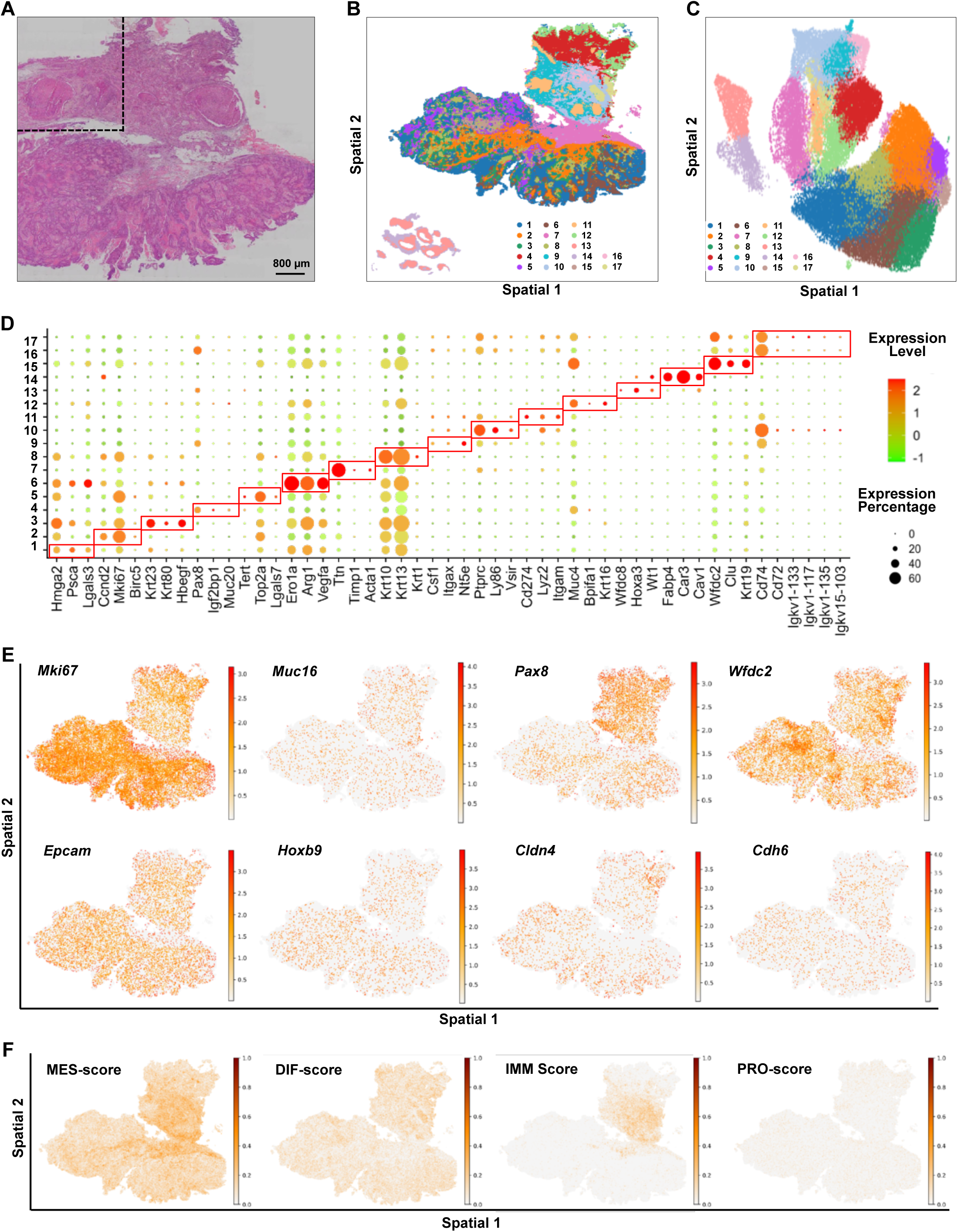
Spatial transcriptomics reveal the cellular landscape of *BTP*-RO tumor at single-cell resolution. A) H&E staining of tumor tissue used for spatial transcriptomics. The region within the dashed outline was excluded due to chip size. Scale bar: 800 µm. B & C) Spatial map and corresponding UMAP reveal 17 spatial clusters across *BTP*-RO tumor and normal control RO tissues (l(45)ower left corner) filtering out low quality regions. The identified regions include: 1: keratinized carcinoma; 2: Fast-growing carcinoma; 3: Cornified carcinoma 1; 4: Pax8^high^ carcinoma; 5: Tert^high^ proliferative carcinoma; 6: Hypoxia-associated carcinoma 1; 7: CAF (cancer-associated fibroblasts); 8: cornified carcinoma 2; 9: Dendritic cells; 10: TAM-1 (tumor associated macrophages, group 1); 11: TAM-2; 12: Secretory carcinoma; 13: RO tubal epithelium; 14: RO stroma; 15: Wfdc2/He4^high^ carcinoma; 16: B cell-1; and 17: B cell-2. D) Dot plot showing top 3 differentially expressed gene markers for each spatial cluster. Dot color and size represent expression level and spot frequency. E) Spatial plots showing expression and localization of canonical HGSC markers *Muc16, Wfdc2 (He4), Epcam, Pax8, Cldn4, Cdh6,* and *Mki67* across the *BTP*-RO tumor section. *Hoxb9* is a ROEC lineage marker. F) Spatial plots showing the expression pattern of signature gene panels defining four molecular subtypes of human HGSC across the *BTP*-RO tumor section. Reference gene panels used for calculating mesenchymal subtype scores of each subtype were extracted from TCGA datasets and presented in the supplement sheet.

### Spatial Transcriptomic Profiling Reveals Necrosis-Induced Cellular Landscape Remodeling

Necrosis has long been associated with enhanced tumor invasiveness across multiple cancers, including high-grade serous carcinoma (HGSC) (66–70). We observed that necrosis represented a defining feature of *BTP*-RO carcinomas (Fig. 3D, 3G, Fig. 4, Fig. 5A). To elucidate the cellular and molecular mechanisms underlying necrosis and its contribution to tumor progression, we performed spatial transcriptomic profiling and stratified the tumor into seven molecularly and morphologically distinct layers: three outer carcinoma layers (OTC1–3), two inner carcinoma layers (INC1–2), a cancer-associated fibroblast (CAF) layer, and the central necrotic region (Fig. 5B-5C). Comparative analyses between outer tumor cells and necrotic core clusters revealed transcriptional gradients linked to hypoxia and necrosis-associated immune signaling (Fig. 5A-5C). Hypoxia-responsive genes *Ndrg1* and *Scel* were enriched in peri-necrotic zones, while *Hif1a* and *Vegfa* displayed spatially restricted expression peaks at the necrosis interface, indicating severe oxygen deprivation and metabolic stress (Fig. 5D).

**Figure 5.**
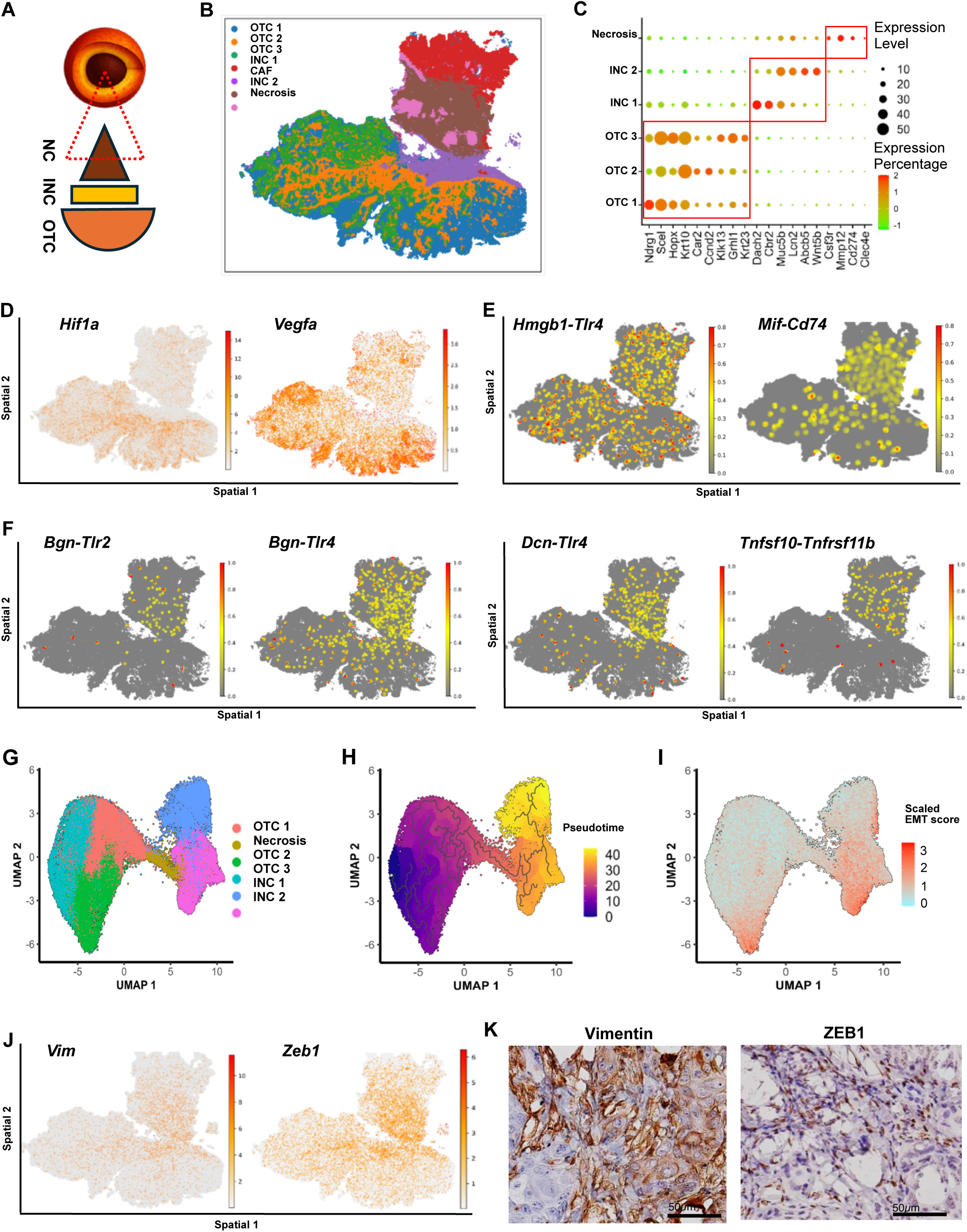
Spatial transcriptomics define necrosis-induced microenvironmental remodeling in *BTP*-RO tumor. A) Schematic representation of tumor architecture, highlighting the central necrotic core (NC), the inner carcinoma layer (INC), and the outer carcinoma layer (OTC). B) Spatially resolved *BTP*-RO tumor regions identified by unsupervised clustering and morphology. Based on both molecular similarity from unsupervised clustering and tumor morphology, tumor tissues were clustered into seven regions, including the outer carcinoma layers 1-3 (OTC1-3), inner carcinoma layers 1-2 (INC1-2), a middle cancer-associated stroma layer (CAF), and the necrotic region. C) Dot plot showing the top 3 biomarkers for tumor region annotation. The dot color and size reflect expression level and frequency of DEGs. D) Spatial map showing expression and location of hypoxia-associated genes such as *Hif1a* and *Vegfa* within the peri-necrotic region (OTC region) of *BTP*-RO tumor tissues. E) Ligand-receptor interaction analysis highlighting expression pattern of genes encoding major components of the alarmin signaling pathways. Presented here are DAMPs (Damage-Associated Molecular Patterns) Hmgb1 and MIF and their respective DAMP-sensing receptors Tlr4 and Cd74 in the peri-necrotic regions and partially necrotic region of *BTP*-RO tumor tissues. F) Ligand-receptor interaction analysis highlighting ligands (*e.g*., *Bgn*, *Dcn*, and *Tnfsf10*, *etc*.) and receptors (Tlr2 and Tlr4, and Tnfsf11b, *etc*.) that trigger signaling pathways governing necrosis-associated inflammation in the necrotic region of *BTP*-RO tumor tissues. G) UMAP restricted to carcinoma/necrosis compartments. H) Pseudo-time trajectory depicting tumor evolution across OTC, INC, and necrotic regions. I) EMT score distribution across OTC, INC, and necrosis clusters showing epithelial-mesenchymal transition gradients. J) Spatial map showing the expression pattern of genes encoding the EMT markers (*Vim*, *Zeb1*) across *BTP*-RO tumor tissues. K) Immunohistochemical analysis validating expression of Vim and Zeb1 (EMT markers) at the protein level. Scale bar: 50 µm.

Ligand–receptor interaction analyses further revealed robust activation of alarmin signaling pathways, with the DAMP molecules *Hmgb1* and *MIF* and their receptors *Tlr4* and *Cd74* concentrated in peri-necrotic regions (Fig. 5E). Additional ligand–receptor pairs, including Biglycan (Bgn), Decorin (Dcn), and Tnfsf10 interacting with Tlr2, Tlr4, and Tnfrsf11b (Fig. 5F), highlighted inflammation-centered networks dominated by Bgn-, Dcn-, and TNF-mediated signaling in necrosis-adjacent zone and necrotic core (71–74), emphasizing necrosis as a key driver of inflammatory propagation.

Pseudotime trajectory analysis mapped necrosis along the tumor’s evolutionary continuum, where outer carcinoma clusters represented pre-necrotic states and inner necrotic cores reflected post-necrotic remodeling (Fig. 5G–H). Spatial proximity to necrosis correlated with increased epithelial–mesenchymal transition (EMT) scores (Fig. 5I), accompanied by elevated Vimentin (Vim) and Zeb1 expression at both transcriptomic and protein levels (Fig. 5J–K). These observations align with established models of ovarian carcinoma progression, wherein necrotic injury promotes EMT, invasion, and metastasis (66–70).

Together, these data reveal that necrosis is not a terminal event but an active organizer of the tumor ecosystem. During *BTP-RO* tumor development, it arises from hypoxia driven by fast-growing HGSC cells, initiates pathogen-free chronic inflammation through DAMP-mediated signaling, promotes EMT, and remodels the microenvironment, thereby driving tumor progression and invasiveness.

### Immune Regulation within the Tumor Microenvironment of BTP-RO Carcinoma

Given that spatial transcriptomic profiling revealed that immune suppression represents a major feature of *BTP*-RO tumors (Figs. 3E-3F and 4F), and necrosis-associated inflammation reshaped the tumor microenvironment, we next examined immune regulation across the tumor’s spatial axis. In the outer carcinoma (OTC) regions, expressions of general immune markers (Ptprc/Cd45, Cd74, H2-Aa, H2-Ab1, *etc.*) were markedly low, defining an immune-poor periphery (Fig. 6A). However, the OTC compartment was enriched in myeloid and M2-like macrophage programs, with *Itgam* and *Csf1r* co-localized with *Arg1* expression, and immunofluorescence confirming strong Arg1-F4/80 overlap (Figs. 6B–D, 6G). These Arg1-high Tumor-associated macrophages suppress T-cell activity through L-arginine depletion (75, 76), and M2 polarization is associated with tumor progression (77–80). Notably, Arg1 intensity correlated with hypoxia (Fig. S4A-4B), reflecting the existence of a hypoxia-driven M2 polarization axis known to promote aggressive tumor behavior and poor prognosis (81). Consistently, spatial mapping of Trac, encoding the T-cell receptor α-chain, showed almost undetectable expression, suggesting limited T-cell infiltration (Fig. 6E). Immunohistochemistry confirmed this pattern, with negligible CD3ε and CD8α staining in OTC regions and sparse T cells in INC zones (Fig. 6F), supporting the presence of an immune-excluded periphery. In contrast, INC regions were not immune-deserted but exhibited checkpoint-mediated suppression. Ligand-receptor (LR) analysis revealed immune checkpoint activation networks involving Cd274-Cd80, Lgals1-Ptprc, Lgals9-Havcr2, and Tgfb1-Tgfbr3 pairs (Fig. 6G), consistent with T-cell exhaustion profiles described in HGSC (82–84). Moreover, co-localization of Igkc, Igka, and Ighm with Lgals9 indicated the accumulation of functionally inactivated B cells at necrotic borders (Figs. 6H–I), consistent with reports that Galectin-9 binding to IgM-BCR attenuates B-cell signaling (85). Finally, high expression of hypoxia-associated genes Hif2a and Ndrg1, together with elevated Slc16a1 and Slc9a1 expression (Figs. 6J–K), demonstrated a hypoxia-driven acidic microenvironment (86). Elevated acidity in tumor tissues is known to impair T-cell cytotoxicity and promote immune tolerance (87–89), providing a biochemical basis for the observed immune suppression.

**Figure 6.**
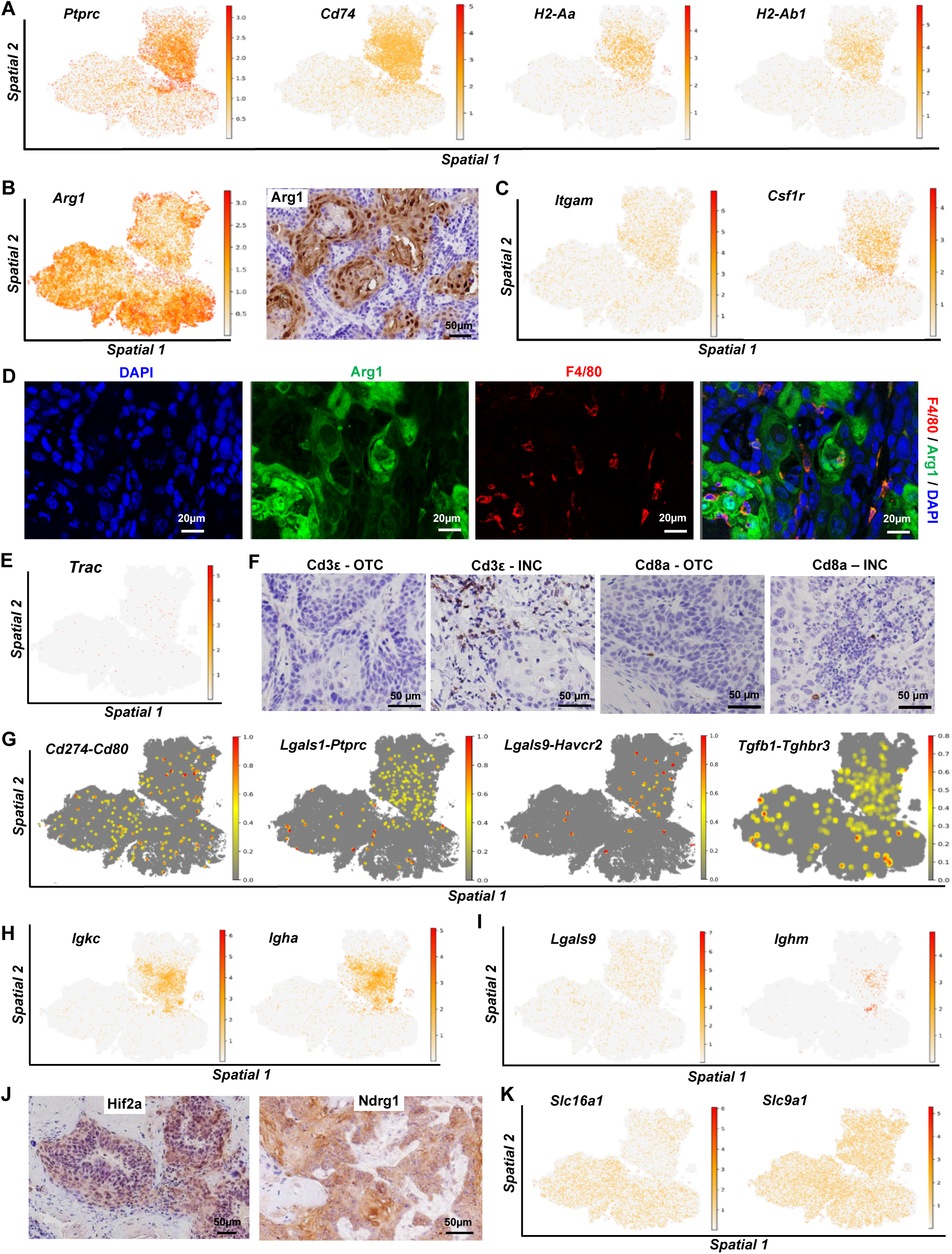
Characterize *BTP*-RO tumor immunosuppressive microenvironment at single cell resolution using spatial transcriptomics. A) Spatial mapping of immune cell biomarkers (*Ptprc*, *Cd74*, *H2-Aa*, *H2-Ab1*) showing immune cell distribution in *BTP*-RO tumor sections. B) Spatial transcriptomics reveals expression pattern of *Arg1* (a classic marker of immunosuppressive microenvironment) in the *BTP*-RO tumor section. On the right is a representative image showing the expression of Arg1 protein. Scale bar: 50µm. C) Spatial map showing *Itgam and Csf1r* (biomarkers of tumor associated macrophage) in the *BTP*-RO tumor tissues. D) Fluorescent IHC validating the colocalization of Arg1 (in green) and F4/80 (in red) proteins (two classic biomarkers of TAMs) in the *BTP*-RO tumor tissues. Nuclei were stained with DAPI (blue). Scale bar: 20 µm. E) spatial map showing expression pattern of *Trac* gene (encoding for the constant region of the T cell receptor alpha chain), indicating minimal T-cell infiltration in the *BTP*-RO tumor tissues. F) Immunohistochemistry showing expression of T cell markers such as Cd3ε and Cd8a protein in the *BTP*-RO tumor. Few Cd3ε+ and Cd8a+ cells confirm spatial transcriptomics results showing sparse T-cell infiltration in *BTP*-RO tumor tissues. Scale bar: 50 µm. G) Ligand-receptor analysis demonstrating enrichment of immunosuppressive cell-cell interactions (Cd274-Cd80, Lgals1-Ptprc, Lgals9-Havcr2, and Tgfb1-Tgfbr3) in the *BTP*-RO tumor tissues. H) Spatial map showing enriched B cell signals in the necrotic zone in the *BTP*-RO tumor. I) Spatial maps showing co-existence of *Lgals9 and Ighm* signaling, evidence for functional inactivation of B cells accumulated in necrosis-adjacent region. J) High expression of *Hif2a* and *Ndrg1* confirm the hypoxic microenvironment in the *BTP*-RO tumor tissues. Scale bar: 50 µm. K) Spatial map showing expression pattern of *Slc16a1* and *Slc9a1,* two acid-base transporters contributing to tumor acidic microenvironment.

Collectively, these data delineate a spatially stratified immunosuppressive architecture in *BTP*-RO carcinoma: an outer immune-excluded, M2-dominated region driven by hypoxia and metabolic suppression; an inner necrosis-associated compartment exhibiting immune exhaustion and checkpoint activation; functionally inactivated B cells near necrosis; and elevated ECM acidity in peri-necrotic zones. Together, these synergistic features establish a profoundly immunosuppressive microenvironment that promotes tumor progression and invasiveness.

## DISCUSSION

Based on the phenotypic, genetic, and developmental resemblance between *BRCA* mutation-associated serous tubal intraepithelial carcinomas arising from fallopian tube epithelial cells and HGSC, HGSC has been regarded as “tubular” in origin, with the fallopian tube recognized as its principal source (20, 21, 23, 24, 40). However, the clinical significance of *BRCA* mutation-associated STICs remains uncertain in the general population, which accounts for 85-90% of all HGSC cases (27–30). The pronounced intra- and intertumoral heterogeneity of HGSC further suggests the possibility of multiple cells of origin. Notably, the female reproductive tract contains additional tubular epithelial systems beyond the fallopian tube. Among them, the rete ovarii represents an anastomosing tubular network derived from mesonephric tubules that invade the developing gonad during embryogenesis and persist as intraovarian, connecting, and extraovarian tubules within the ovarian hilus (36, 90). Historically regarded as a vestigial mesonephric remnant, the RO has been largely overlooked, partly due to its small size, deep hilar location, and limited tissue availability in humans. The lack of suitable animal models has also hindered functional studies (35, 91, 92). Recent high-resolution molecular profiling and lineage-tracing analyses, however, have redefined the RO as a structurally heterogeneous and transcriptionally active organ (34, 45). The extraovarian compartment exhibits PAX8 expression, ciliation, and secretory machinery, suggesting a potential paracrine or “antenna-like” role in ovarian physiology (34). Morphologically, RO epithelial cells share features with fallopian tube secretory and ciliated cells, including columnar architecture and partial overlap in immunophenotypic markers, yet maintain distinct transcriptional identities compared to fallopian tube or endometriotic epithelium (93). In the present study, we demonstrate that ROECs can undergo malignant transformation upon manipulation of canonical ovarian cancer oncogenes and tumor suppressors, supporting the RO epithelium as a previously unrecognized tubular source contributing to pelvic and ovarian HGSC. Our results identify ROECs as a previously unrecognized cell of origin for ovarian high-grade serous carcinoma. Because rupture of RO cysts could release transformed cells into the pelvic cavity, ROECs may also serve as a source of disseminated pelvic HGSC. These findings expand the current understanding of the cellular origins and pathogenesis of pelvic and ovarian HGSC. Moreover, the cellular and molecular targets identified in this study provide a foundation for developing origin-informed strategies for prevention and early detection of RO-derived HGSC, as well as for discovering targeted therapeutics to improve overall survival in patients with HGSC.

A limitation of the present study is that these findings are derived from mouse models. However, their clinical relevance is supported by historical clinicopathologic reports documenting benign and, albeit rarely, malignant neoplasms arising from RO epithelium (94–96). The rarity with which RO-associated HGSC is recognized in human likely reflects two confounding factors: first, most HGSCs are diagnosed at advanced, disseminated stages that obscure the primary site; and second, the RO’s close anatomic and structural continuity with both the ovary and the fimbriae region of the fallopian tube complicates accurate origin assignment. Nevertheless, several lines of phenotypic similarity from our histologic analyses and molecular concordance from spatial transcriptomic profiling strengthen the biological link between RO-derived neoplasia and human HGSC. Human HGSCs frequently exhibit tubal features and can arise from fallopian tube epithelium, while the RO, particularly its extraovarian compartment, contains both secretory and ciliated cells, a cellular composition permissive for serous (tubular-type) transformation (35, 36, 93, 94). Moreover, the brisk proliferation and tumor necrosis characteristic of HGSC were faithfully recapitulated in our RO-derived tumors (97, 98). At the tumor microenvironmental level, HGSCs, especially those classified to the TCGA/consensus “mesenchymal” molecular subtype, display pronounced stromal activation and immunosuppressive features, consistent with the immune-suppressive microenvironment observed in *BTP*-ROEC-derived tumors (46, 65, 99, 100). Finally, overexpression of canonical HGSC markers in our model, including MUC16/CA125, WFDC2/HE4, PAX8, and KRT7, mirrors the molecular profiles of human HGSC, further underscoring its translational relevance (59, 101–104). Collectively, these findings suggest that dysregulated RO tubules may plausibly give rise to a subset of human HGSCs and warrant targeted lineage-tracing and comparative transcriptomic studies to determine the extent to which clinically annotated HGSCs originate from the RO.

Interestingly, recent evidence suggests that both ovarian and fallopian tube epithelia can contribute to HGSC development (21, 33). Because HGSC exhibits “tubular” characteristics such as PAX8 expression, ciliated and secretory differentiation, and transcriptional similarity to fallopian tube secretory cells, the presence of these features in ostensibly ovarian-origin tumors has been long puzzling, given that the ovarian surface epithelium (OSE) is mesothelial rather than Müllerian in nature (105, 106). Our findings, together with prior reports, identify rete ovarii epithelial cells as a distinct intraovarian population of PAX8-positive, ciliated, and secretory epithelial cells within intraovarian and collecting rete tubules (34, 35, 107), providing a compelling explanation for this paradox. Developmental and histopathologic analyses demonstrate that these mesonephric-derived tubules persist into adulthood and can give rise to intraovarian cystic or proliferative lesions (92, 108, 109). This raises the possibility that a subset of ovarian inclusion cysts, traditionally attributed to OSE invagination or metaplasia, may instead arise from rete-derived epithelium (32) (110). Given their molecular similarity to fallopian tube secretory cells, ROECs may undergo analogous oncogenic transformation upon acquisition of TP53, BRCA1/2, or PTEN pathway alterations, leading to the formation of inclusion-type cystic lesions and invasive HGSC This model reconciles the occurrence of “ovarian” HGSCs displaying Müllerian molecular signatures in the absence of identifiable fallopian tube precursors. Collectively, these findings support a revised paradigm in which a subset of inclusion cysts originates from ROECs, thereby revealing novel cellular and molecular targets for prophylactic prevention, early detection, and therapeutic intervention.

In summary, our findings demonstrate that epithelial cells within the unconventional tubular network of the rete ovarii can undergo malignant transformation, giving rise to high-grade serous carcinomas with histologic and molecular features closely recapitulating human HGSC. We propose that the rete ovarii epithelium, a persistent tubular structure within the ovarian hilus, constitutes an overlooked cell source contributing to the development of a subset of HGSC, thereby expanding the spectrum of potential precursor populations beyond the fallopian tube. This RO-based HGSC model provides a conceptual and experimental framework for developing origin-informed strategies aimed at effective prevention, early detection, and targeted treatment of RO-derived HGSC.

## MATERIALS AND METHODS

All experimental procedures were conducted in accordance with institutional and sponsor regulations. No human subjects were involved in this study. All animal procedures, including mouse handling, breeding, and treatment, were approved by the Institutional Animal Care and Use Committee (IACUC) of Massachusetts General Hospital (MGH).

### Establishment of RO Tubal Epithelial Cell Lines

Rete ovarii tubal epithelial cells were isolated from BT-Cas9 mice. Three-dimensional (3D) organoid cultures were established using Matrigel Matrix (CLS354234; Corning, Glendale, AZ) according to the manufacturer’s protocol. Cells were also maintained as two-dimensional (2D) monolayers in cell culture dishes. All cultures were grown in Advanced DMEM-F12 (11320033; Gibco, Thermo Fisher Scientific, Waltham, MA) supplemented with B27, EGF, and a TGFBR1 inhibitor following a previously published formulation (111). To enrich epithelial cells, the mixed cell population was sorted using CD326 (EpCAM) MicroBeads (130-105-958; Miltenyi Biotec, Auburn, CA), selecting for CD326-positive cells.

Cells were transduced with human Adenovirus-CRE Type 5 (dE1/E3) containing a CMV promoter-driven RFP reporter (1770; Vector Biolabs, Malvern, PA) to activate recombination at loxP sites, following the approved MGH protocol (PIBC Registration #2017B000085). Gene editing was performed using the Gene Pulser Xcell Electroporation System (Bio-Rad Laboratories, Hercules, CA) to introduce *Brca1/2* and *Pten* sgRNAs. *Brca1-Trp53-Pten* and *Brca2-Trp53-Pten* engineered cells were mixed in equal proportions and injected into immunocompetent recipient mice with similar genetic background.

### Establishment of BTP Syngeneic Tumor Models

Transgenic mice carrying heterozygous *Trp53* knockout and homozygous floxed *Brca* alleles were purchased from The Jackson Laboratory (Strain #012620; Bar Harbor, ME, USA). Cas9-expressing mice (Strain #026179; The Jackson Laboratory) were also obtained and subsequently bred in-house to establish the BT-Cas9 line. *Pten* and *Brca* knockouts were introduced using the CRISPR-Cas9 system.

To establish syngeneic tumor models, genetically modified RO epithelial cells were injected intraperitoneally (IP) into 8-week-old female mice with matching genetic backgrounds (n = 6 for immunocompetent mice; n = 8 for NSG mice). Each mouse received approximately 1.5 × 10⁶ cells and 50µg tacrolimus (F-4900; LC Laboratories, Woburn, MA). Mice were monitored daily and euthanized after approximately six months to assess tumorigenesis.

### Immunohistochemistry and Immunofluorescence

Immunohistochemistry (IHC) was performed to evaluate the expression of high-grade serous ovarian carcinoma (HGSC) biomarkers and immune cell signature molecules. Tissues were fixed in 4% paraformaldehyde, embedded in paraffin, and sectioned for analysis. After deparaffinization, slides were blocked with 5% normal donkey serum and incubated with primary antibodies at 4 °C for 16 h. IHC signals were visualized using a polymer-based detection system and a DAB assay kit (Vector Laboratories, Burlingame, CA) as previously described (112). Immunofluorescence staining was performed according to established laboratory protocols (113) to assess protein localization. Secondary antibodies included anti-rat IgG (4418; Cell Signaling Technology, Danvers, MA) and anti-rabbit IgG (39572; Cell Signaling Technology).

### Stereo-seq Sample Preparation and Sequencing

Two paraffin-embedded tissue sections (1 cm × 1 cm each) were mounted onto Stereo-seq Chip N slides (STOmics, San Jose, CA, USA), comprising one tumor specimen and one normal fallopian tube (FT) block containing multiple FT segments. Sections (5 µm thick) were stained using a fluorescently labeled single-stranded DNA (ssDNA) protocol following the manufacturer’s instructions (Qubit™, Invitrogen, Waltham, MA, USA). Confocal imaging for nuclear staining quality control was performed on a Zeiss Axio Observer.Z1 microscope (Zeiss, Oberkochen, Germany).

cDNA synthesis was conducted using the Stereo-seq Transcriptomics N Kit v1.0 (STOmics). Library preparation followed the DNA NanoBall (DNB) approach using the Stereo-seq 16 Barcode Library Preparation Kit v1.0 (STOmics). Approximately 20 ng of cDNA was used per library, purified with SPRIselect beads (Beckman Coulter, Brea, CA, USA), and sequenced on the DNBSEQ-T7 platform (Complete Genomics, San Jose, CA, USA). Each DNB contained a 25 bp cell identifier (CID), a 6 bp molecular identifier (MID), and a 10 bp sample barcode. Library quality was assessed using an Agilent 2100 Bioanalyzer (Agilent Technologies, Santa Clara, CA) with a high-sensitivity DNA chip.

### Stereo-seq Data Processing

FASTQ files from the DNBSEQ-T7 platform were processed using the Stereo-seq Analysis Workflow (SAW) count pipeline on the ERISTwo HPC cluster at MGH. The reference genome was generated using the SAW makeRef pipeline based on the GRCm39 mouse genome, followed by alignment using the STAR algorithm.

Low-quality reads (MID base quality < Q10 or > 1 ambiguous base), unmapped reads, low-mapping-quality reads (MAPQ < 255), and rRNA-derived reads were excluded. Cell segmentation (cellbin annotation) was guided by ssDNA images, assigning reads to cells based on image intensity and morphology.

Dimensionality reduction was performed by principal component analysis (PCA), and clustering was conducted using the Leiden algorithm. Cell bins with < 100 unique molecular counts and genes expressed in < 10 cells were removed using the *Scanpy* package. Differential gene expression between clusters were assessed with the Wilcoxon rank-sum test. Cells with > 10% mitochondrial gene content were excluded. Spatial visualization of cell bins and bin20 distributions was generated using StereoMap (STOmics).

### RNA Sequencing and Data Analysis

Total RNA was extracted directly from cultured cell lines. High-quality sequencing libraries were prepared and sequenced on an Illumina NovaSeq 6000 system. The reference genome (GRCm39) was indexed using BWA and Samtools, and reads were aligned to the reference with BWA-MEM (v0.7.17) using default parameters and read group information. The resulting SAM files were converted to BAM format for downstream processing.

Differentially expressed genes (DEGs) were identified using edgeR (v3.36.0) in R (v4.4.2). Gene set enrichment analysis (GSEA) was performed using GSEA software (v4.3.3; Broad Institute, MA, USA) with datasets from the MSigDB database (114, 115). Pathways with nominal *p*-values (NOM *p*-val) and false discovery rate *q*-values (FDR *q*-val) > 0.05 were excluded from further analysis. Enrichment plots were generated using the GSEA software, and summary visualizations were produced with the ggplot2 package in R.

### Single-Cell Transcriptomics Preparation

Tumor samples were generated by pooling tissues from five mice (n = 5), yielding approximately 1 × 10⁶ cells to construct a single-cell RNA sequencing (scRNA-seq) library targeting 10,000 cells, as described previously (112). Control samples were prepared similarly using tissues from five mice (n = 5).

Tissues were finely minced into ∼1 mm² fragments on ice, rinsed with cold PBS, and digested at 37 °C for 40 min with gentle agitation (150 rpm) in an air-bath shaker. The enzymatic digestion buffer contained 450 U/mL collagenase I, 150 U/mL collagenase II, 450 U/mL collagenase III, 450 U/mL collagenase IV, 0.8 U/mL elastase, 300 U/mL hyaluronidase, 250 U/mL DNase I, and 1 U/mL dispase. Following digestion, cells were diluted in cold RPMI 1640 medium containing 2% fetal bovine serum (FBS), filtered sequentially through 70 μm and 40 μm strainers, and centrifuged at 1,500 rpm for 5 min at 4 °C. The pellet was treated with ACK lysis buffer for 10 min at room temperature to remove red blood cells, washed, and resuspended in ice-cold PBS with 0.04% bovine serum albumin (BSA). Cell viability was assessed using a TC20™ Automated Cell Counter (Bio-Rad, Hercules, CA, USA) prior to library construction.

Single-cell RNA-seq libraries were generated using the 10x Genomics Chromium Next GEM Automated Single Cell 3′ Reagent Kit v3.1, which uses poly(dT) primers to capture polyadenylated mRNA transcripts. Cell suspensions were loaded onto a Chromium Single Cell Chip together with the reverse transcription master mix and single-cell 3′ gel beads, targeting approximately 10,000 cells per channel. Reverse transcription and barcoding were performed on a C1000 Touch Thermal Cycler (Bio-Rad). cDNA quality and quantity were assessed using an Agilent 2100 Bioanalyzer (Agilent Technologies, Santa Clara, CA, USA) with a high-sensitivity DNA chip. Gene expression libraries were constructed by amplifying 50 ng of cDNA through 14 PCR cycles, and final libraries were sequenced on an Illumina HiSeq 2500 platform following the manufacturer’s recommended parameters.

### Bioinformatics and Transcriptomics Analysis

Raw scRNA-seq FASTQ files were processed using Cell Ranger (v9.0.0; 10x Genomics) with the single-cell 3′ v3 chemistry and a reference genome based on GRCm39, built using the Cell Ranger *mkref* pipeline. Downstream analyses were performed using the Seurat package (v5.2) (116). Low-quality cells with insufficient RNA counts were removed. Specifically, cells with <300 or >5,000 detected features, or >10% mitochondrial gene content, were excluded. To eliminate erythrocyte contamination, cells expressing >10% of *Hba-a1*, *Hba-a2*, *Hbb-bs*, or *Hbb-bt* transcripts were also filtered out.

Dimensionality reduction was performed by principal component analysis (PCA), and the first 15 principal components were used for unsupervised clustering with the Louvain algorithm. Clusters were visualized using Uniform Manifold Approximation and Projection (UMAP). Two single-cell datasets were merged into a unified atlas by aligning gene expression matrices and applying SCTransform normalization. Cell types were manually annotated based on differentially expressed genes identified by the Wilcoxon rank-sum test.

Spatial transcriptomics data were analyzed using Scanpy (v1.10.4; Python v3.12.3), Seurat (v5.2), and Monocle 3 (117–119) for pseudotime trajectory reconstruction. Cells with fewer than 2,000 detected counts and genes expressed in fewer than 500 cells were removed. Data were normalized for sequencing depth using library size normalization followed by logarithmic transformation.

Differential expression analysis comparing cell-type marker genes across spatial clusters was conducted using the Wilcoxon rank-sum test. Marker gene expression patterns were visualized with standard Scanpy plotting functions, including violin plots, dot plots, and spatial feature maps, to illustrate intratumoral heterogeneity. Carcinoma cell markers were compared with published high-grade serous ovarian carcinoma (HGSC) markers (120) and immune-suppressive cell markers (111, 121).

Cell-cell communication (CCC) analysis was performed using LIANA (v1.5.1) (122). To evaluate EMT and hypoxia within the spatial transcriptomics data, gene sets *HALLMARK HYPOXIA* and *HALLMARK EPITHELIAL MESENCHYMAL TRANSITION* were retrieved from MSigDB. Gene set activity scores were calculated for each cell using the UCell package (v2.1.2) (123).

## ACKNOWLEDGEMENT

This work was partially supported by the National Cancer Institute/National Institute of Health (2R01CA197976, 1R01CA201500, 5R01CA279385, CW), the Colleen’s Dream Foundation (no number, CW), the Ruggles Family Foundation (CW), the Vincent Memorial Hospital Foundation (CW), and the Vincent Department of Obstetrics and Gynecology, Massachusetts General Hospital.

## DATA AVAILABILITY

Raw and processed sequencing data for bulk RNA-seq, single cell RNA-seq, and spatial transcriptomics are available at Series: GSE306573, GSE305806 and GSE305977. Any additional information required to reanalyze the data reported in this paper is available from the lead contact upon request.

## AUTHOR CONTRIBUTATION

C.W. acquired the funding and supervised this project; P.C., C.W. designed the research; P.C., X.Z., J.R. performed research; X.Z., performed bioinformatics analyses/visualization; X.Z. and J.R. conducted spatial transcriptomics; P.C., X.Z., and J.M. contributed to SC-RNAseq data collection and analyses. P.C., X.Z., J.R., X.L., J.L., H.W., D.S., M-L.M., B.C., G.K.,O-A.A, performed cell culture, sample collection, and histological studies. P.C., J.R., X.L., J.L., H.W., D.S., M-L.M., B.C., G.K., O-A.A, J.M., and J-S.D contributed to manuscript review and editing; X.Z., and C.W. wrote the manuscript.

## Notes

Disclosures: The authors declare that they have no conflict of interest

https://www.ncbi.nlm.nih.gov/geo/query/acc.cgi?acc=GSE305806

https://www.ncbi.nlm.nih.gov/geo/query/acc.cgi?acc=GSE305977

https://www.ncbi.nlm.nih.gov/geo/query/acc.cgi?acc=GSE306573

### Competing Interest Statement

The authors have declared no competing interest.

