## supplementary information for "Rete Ovarii Epithelial Cells as an Unappreciated Cell of Origin for Pelvic and Ovarian High-Grade Serous Carcinoma"

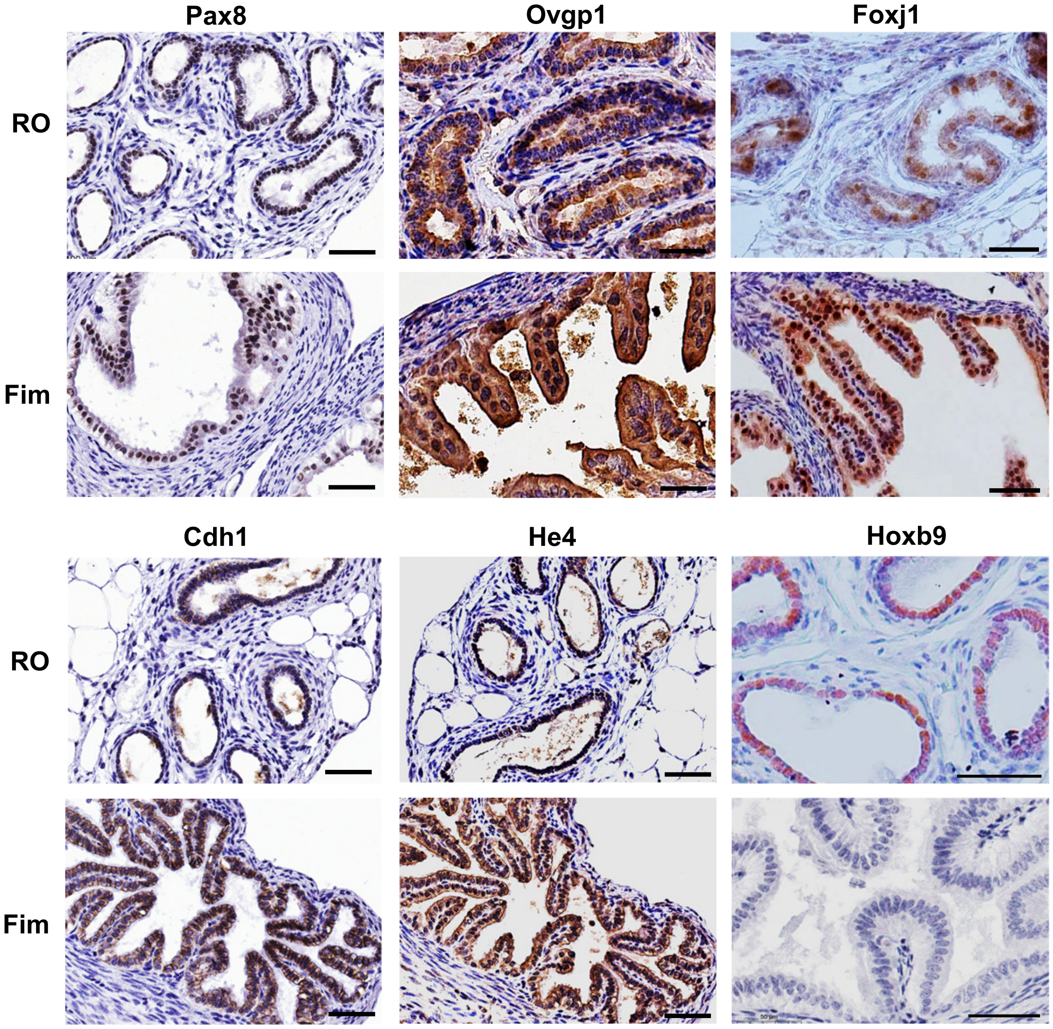


**Figure S1. Representative images showing that RO and FT epithelia share histologic and immunophenotypic features.** The expression of common fallopian tube epithelial cell biomarkers such as Pax8, Ovgp1, Foxj1, Cdh1 (E-cadherin), and Wfdc2 (He4) are highly expressed in both fallopian tube and rete ovarii epithelial cells. Scale bar: 50μm.

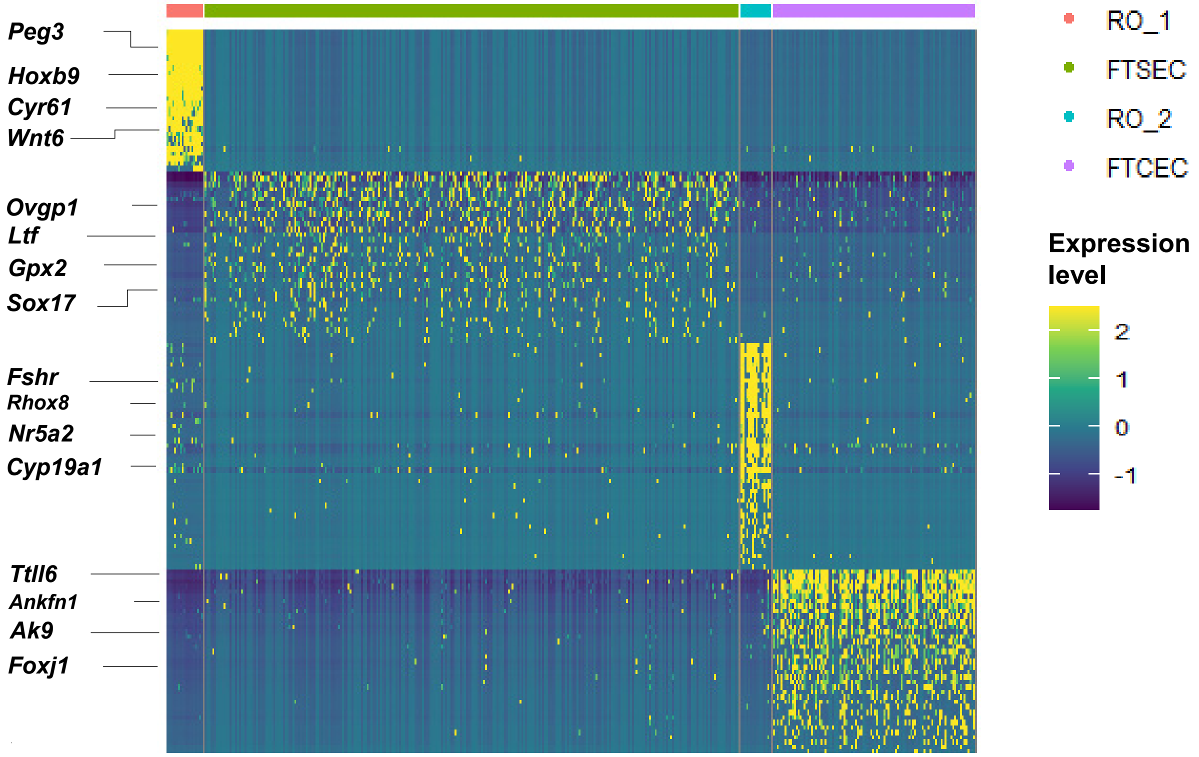


**Figure S2, Heat map showing differentially expressed genes in Rete ovarii (RO) and fallopian tube (FT) epithelial cells.** Unsupervised clustering based on single cell RNA sequencing data re-clustered RO epithelial cells into *Hoxb9* and *Peg3* high cluster 1 (RO_1) and *Fshr* and *Nr5a2* high cluster 2 (RO_2). The FT cells were also divided two subclusters based on their gene expression pattern.

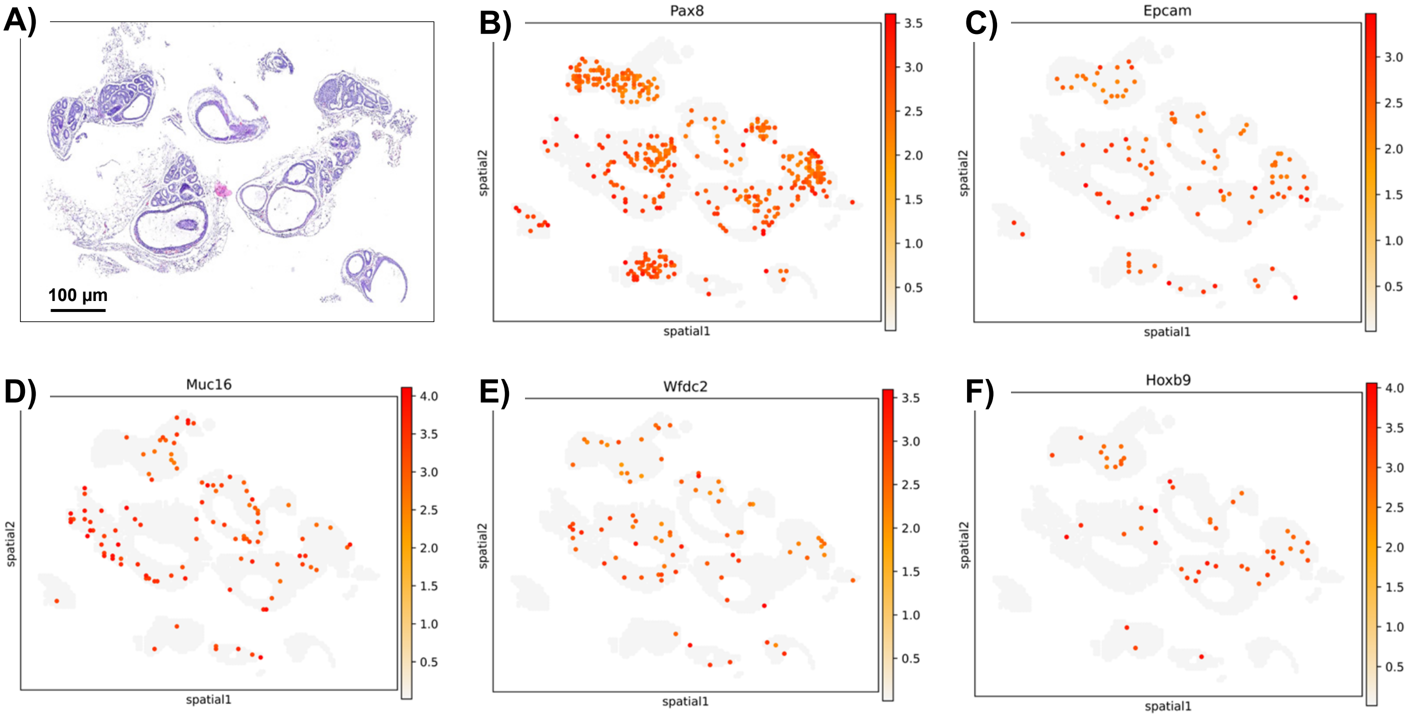


**Figure S3. spatial mapping *Pax8*, *Epcam*, *Muc16, Wfdc2,* and *Hoxb9* in normal rete ovarii (RO) tubes.** A) Histology of isolated normal RO tubes (H-E staining). Scale bar: 100µm. B-F) Expression pattern of *Pax8*, *Epcam*, *Muc16* (encoding for CA125 peptide), *Wfdc2* (Encoding for He4 protein) and *Hoxb9* in normal rete ovarii tubes. Please note that general fallopian tube epithelial cell markers are also expressed in normal RO control tissue detected by spatial transcriptomics.

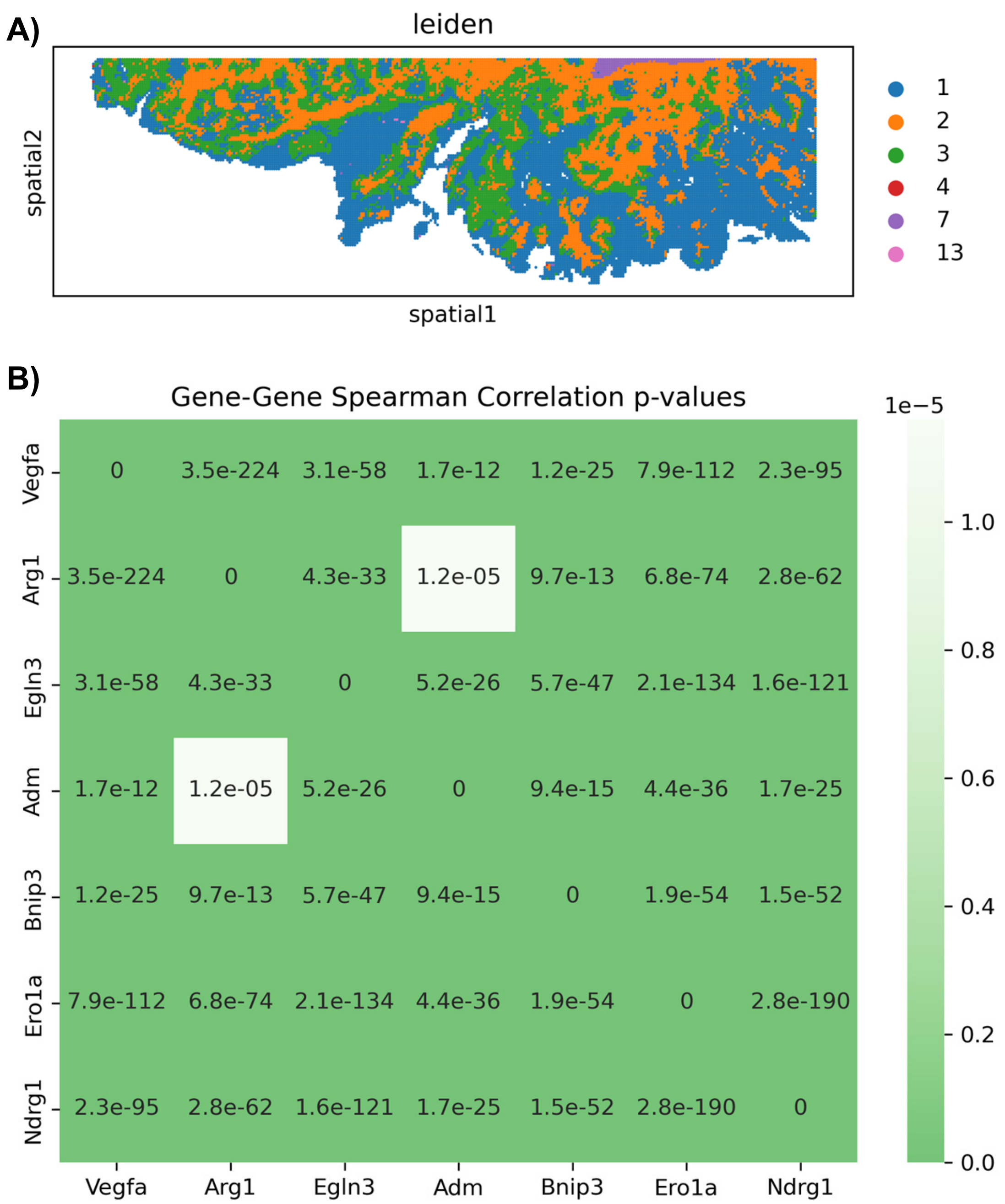


**Figure S4. Arg1 intensity correlates with hypoxia in the peri-necrotic zone.** (A) Peri-necrotic zone selected for analyzing the gene expression correlation between Arg1 and genes encoding for classic hypoxia sensors in tumor tissues. (B) Correlation matrix showing *P* values for the expression relationships between Arg1 gene and classical Hif1a-associated hypoxia sensor genes (*Vegfa*, *Egln3*, *Adm*, *Bnip3*, *Ero1a*, and *Ndrg1*) in the peri-necrotic zones of BTP-RO tumor tissues. Statistical significance was determined using the Spearman rank correlation test.
